# Accurate imputation of inversions in human genomes using different algorithms and data sources

**DOI:** 10.64898/2026.01.23.701363

**Authors:** Illya Yakymenko, Adrià Mompart, Mario Cáceres

## Abstract

Complex genomic regions harbor different structural arrangements that can mutate quite rapidly, which makes determining their functional effects very difficult. Characterization of inversions originated by homologous mechanisms is especially challenging due to the presence of inverted repeats at the breakpoints and the fact that most of them are recurrent. Imputation can infer missing genotypes, but it has been mainly limited to simple variants and little is known about how well it works for human inversions. Here, we tested five common imputation programs to impute a set of 52 inversions experimentally genotyped in multiple samples that lacked SNPs in perfect linkage disequilibrium. Using whole genome sequencing data and simulated microarrays with variable SNP density, we found that 40.4-75.5% of inversions could be accurately imputed in three human populations by at least one program, with results depending mainly on the number of SNPs available, the genotyped samples and the recurrence of inversions. Also, genotype probability filtering was a key factor for inversion imputation accuracy. In particular, Minimac4 and IMPUTE5 showed more accurately imputed inversions and less poorly imputed individuals with respect to the other methods. This work therefore contributes to optimize inversion imputation, making possible the study of their functional impact.

## Introduction

During the last decades there has been a great interest in the complete characterization of human genomic variation and its association with phenotypic traits and disease susceptibility. Traditionally, SNPs and small indels have been the main target, with many large-scale projects across multiple individuals of diverse populations, such as the 1000 Genome Project (1KGP) (1, 2) or the Genome Aggregation Database (gnomAD) (3). In addition, several studies using diverse genomic techniques are finally allowing us to identify all the other more-complex structural variants (SVs) (2, 4–8). However, due to the technical challenges of the detection of these variants, in most cases there are just a small number of reliable genotypes, which complicates determining their functional impact.

Specifically, inversions are a type of SVs that just reverse the orientation of a DNA segment. This confers inversions unique biological properties and they have been associated to phenotypic traits and adaptation in multiple species, but it also makes them particularly difficult to study (9). Moreover, while inversions generated by non-homologous mechanisms typically have nearby variants in perfect linkage disequilibrium (LD) that facilitate to estimate accurately the inversion status (10), inversions mediated by highly identical inverted repeats (IRs) often undergo recurrent toggling between the original and inverted orientations via non-allelic homologous recombination (NAHR) (10–12). This dynamic process increases haplotype diversity and, as a consequence, there is no single variant or set of variants in high LD with the inversion that can reliably predict its orientation. Therefore, low-throughput experimental techniques are required to determine the genotype of the inversions, and their effects have been likely missed in existing genome-wide associated studies (GWAS), which are based mainly on SNPs (9, 10).

GWAS is the standard procedure to check the association of genetic variants to phenotypic traits in large cohorts. These analyses were performed extensively with microarray data, that allow a cost-efficient genotyping of a limited amount of common SNPs in high number of samples, and now they are done increasingly by whole-genome sequencing (WGS) at lower or deeper coverage (13, 14). Imputation is widely used to infer missing variant genotypes for downstream analyses, and multiple imputation algorithms have been developed, each one with its strengths and weaknesses (15–17). Some of the most popular tools are Beagle, which uses a haplotype cluster graph model (18) and IMPUTE2, that applies a Markov chain Monte Carlo (MCMC) framework, which scales poorly (19). Moreover, there are other improved methods, such as IMPUTE5, that enhances this with positional Burrows-Wheeler transform (PBWT) for fast, accurate imputation from large panels (20), or Minimac4, which employs a space state reduction while using a Monte-Carlo chain haplotyping (MaCH) procedure (21). In addition, scoreInvHap was designed to infer genotypes of a few inversions by creating haplotype clusters with flanking SNP information and it has been used to examine their phenotypic associations (22). However, there is still very little information on the performance of different imputation methods for SVs, given the specific characteristics of these variants (23, 24).

In this study, we took advantage of the generation of a valuable data set of experimentally validated genotypes of 134 inversions in 95-550 diverse samples (10, 11) to evaluate the performance of human inversion imputation by benchmarking five widely used imputation tools, with a particular focus on those inversions lacking variants in perfect LD. Our primary aim was to assess the best strategy to impute inversions using current state-of-the-art methods and the main factors affecting imputation reliability, in order to be able to determine their functional effects.

## Materials and Methods

### Small variant and inversion data

All analyses were carried out using the GRCh38 (hg38) human genome reference assembly. The small variant information (including SNPs and small indels), population of origin and sex of each individual for the reference and target panels used for phasing and imputation were extracted in VCF format from the 1KGP high-coverage WGS dataset (1KGP-HC), which can be accessed at https://ftp.1000genomes.ebi.ac.uk/vol1/ftp/data_collections/1000G_2504_high_coverage/working/20220422_3202_phased_SNV_INDEL_SV/ (2). The genetic maps required for phasing and imputation were obtained at https://bochet.gcc.biostat.washington.edu/beagle/genetic_maps/. VCFs files were filtered to contain only biallelic SNPs and exclude singletons using BFCtools 1.19 (with arguments *view--min-ac 2:minor -M2 -m2 -v snps*) (25). No additional minor allele frequency (MAF) filter was applied.

Highly-accurate experimental inversion genotypes to carry out and validate the imputation procedure were obtained in previous studies by different PCR-based methods (10, 11). The dataset includes 52 inversions without known tag-SNPs in perfect LD (*r*^2^ = 1) across African (AFR), European (EUR) and East-Asian (EAS) super-populations based on a previous analysis of the original set of 134 inversions with 1KGP-HC data (Supplementary Table 1). Inversions had been genotyped in a variable number of 86-438 AFR, EUR and EAS 1KGP individuals depending on the technique used (Supplementary Fig. 1). Singleton inversions with less than two variant alleles in the genotyped individuals across populations were not considered in the analysis.

### Variant phasing

Most imputation methods, except for scoreInvHap, use haplotype information to impute unknown variants and they require the input data to be phased. Thus, we rephased the 1KGP-HC VCF files to assign each of the experimental inversion genotypes to the corresponding haplotype. For this, we extracted the variants of an extended region including the inversion and up to 200 kb upstream and downstream from its breakpoints (Supplementary Table 1) with BCFtools (25). Available inversion genotypes were added as another variant (0/0, 0/1 or 1/1) in the VCF in the central position of the inversion region, using bash and vcf-sort (VCFtools 0.1.16) (25). Males were treated as diploid for chromosome X, transforming hemizygous 0 and 1 genotypes to 0/0 and 1/1. Next, we tested phasing with Beagle v5.4 (22Jul22.46e) and SHAPEIT5 (5.1.1) (18, 26) and the results were compared using a Wilcoxon signed-rank test. For Beagle, phasing has been conducted using default parameter settings. For SHAPEIT5, the *phase_common* and the *phase_rare* steps were run respectively with the arguments *--filter-maf 0.05*, and *--pbwt-modulo 0.005*. Then, the processed VCFs were divided by the super-populations using BCFtools (25).

### Inversion imputation

To benchmark inversion imputation performance we used Beagle 5.4 (22Jul22.46e), Impute2 (2.3.2), Impute5 (1.2.0), Minimac4 (4.1.6) and scoreInvHap (1.30.0) (18–22). All programs were run independently for different super-populations with default parameters, specifying human effective population size (N_e_) as 20,000 when possible and adding *--pbwt-cm 0.005* to Impute5. To determine imputation accuracy of most programs, a leave-one-out (LOO) strategy was employed. In this approach, an individual with masked inversion genotype is the target for the imputation, using the rest as the reference data, and the process is repeated for all the individuals of the super-population. Imputation accuracy of each method for the different inversions and super-populations was determined by comparing all imputed and experimentally obtained inversion genotypes with the LD metric *r*^2^ using PLINK (1.90b7.2), with parameters *--ld-window-kb 1000000 --ld-window 999999 --ld-window-r2 0* (27). The imputation process was repeated five times for each method with different random seeds, and the median *r*² value of the five replicates was taken as imputation accuracy. For scoreInvHap (22), the input LD values (*r*^2^) of the inversion with small variants in the 200 kb extended region were calculated from the VCFs of each super-population with PLINK (27), using the same parameters as before. The remaining input information was obtained processing the data according to scoreInvHap documentation (https://bioconductor.org/packages/release/bioc/html/scoreInvHap.html). As scoreInvHap is just based on the LD values and does not consider the real inversion genotypes, imputation accuracy was obtained directly by calculating the *r*^2^ between experimental and imputed genotypes in each super-population, without requiring LOO processing.

In addition, to check if in some cases imputation improves when increasing the number of available genotypes, imputation for all methods except scoreInvHap was repeated for the global (GLB) population, combining together the AFR, EAS and EUR individuals. The imputation, LOO and *r*² calculation was done as previously described by using as reference the unsplit VCF files. All imputation results are provided in Supplementary Table 2.

### Genotype probability filtering

For each imputed variant, imputation algorithms report genotype probabilities (GP), which a value of 0 to 1, representing the likelihood of the three possible genotypes (reference homozygote, heterozygote and alternative homozygote, respectively). In all analyses, the inversion genotype with the highest GP was selected as the most probable one, and, for simplicity, GP refers to the maximum GP observed for each inversion and individual. GP values were extracted from the GT (GT:GP) field for the imputed variants of the VCF file. To determine the effect on imputation accuracy of eliminating low reliable imputed genotypes, different GP filters, ranging from 0 to 0.95, were applied to remove those individual genotypes that did not meet the minimum GP criterion prior to the calculation of the *r*² in the LOO process. Alternatively, to compare the effect of filtering the same number of low reliable imputed genotypes across programs, we also removed 5% to 25% of samples with lowest GP for a given inversion and imputation method prior to *r*² calculation. Ties with the same GP values for an inversion were resolved by selecting randomly the filtered samples.

All follow-up comparisons, except for when specifically mentioned, have been carried out applying a GP filtering threshold of ≥0.9. Imputable inversions in a population were considered those with an imputation accuracy of *r*^2^ ≥ 0.8 and more than 50% remaining samples after the 0.9 GP filtering.

### Imputation in SNP arrays

To check inversion imputation performance from SNP genotyping array data, two representative SNP arrays with different density have been simulated: the high-density Illumina Infinium Omni 2.5-8 v1.3 array (Omni), interrogating approximately 2.37 million SNPs; and the low-density Illumina Infinium Global Screening Array v2.0 (GSA), with approximately 650,000 SNPs. To simulate these arrays, CSV files containing the SNP information in the GRCh38 assembly were downloaded from https://support.illumina.com/array/array_kits/humanomni2_5-8_beadchip_kit/downloads.html and https://support.illumina.com/array/array_kits/infinium-global-screening-array/downloads.html. Then, the position of the array SNPs was retrieved and those within the inversion plus 200 kb flanking region were selected from the filtered 1KGP-HC VCFs using BCFtool. (25). Finally, splitting samples per population, imputation, LOO and calculation of imputation accuracy were done as described above (Supplementary Table 2).

### Statistics and linear model analysis

Comparisons between phasing and imputation methods or reference panels were performed using a non-parametric paired Wilcoxon signed-rank test, including always the set of inversions in common to the conditions compared. Comparison between populations were done with a non-parametric unpaired Wilcoxon rank-sum test, since the inversions polymorphic in each population are not the same. A regular Person correlation coefficient (*R*) was used to compute correlations between programs or variables. Linear regression models have been fitted to the imputation accuracy (*r*² ) using different biological features as independent variables, such as number of genotyped individuals, ratio between IR and inversion length, MAF, number of SNPs in the region and presence in chromosome X (Supplementary Table 3), with the formula *r*² ∼ Predictors in R. Samples from all three populations have been combined and, with the exception of scoreInvHap, one linear model per imputation method has been fitted with and without GP < 0.9 filtering.

## Results

### Comparison of inversion imputation methods

As already mentioned, we took advantage of the highly-accurate inversion genotypes from multiple 1KGP individuals (10, 11), together with their high-coverage WGS data (2), to evaluate how well different methods and imputation strategies infer the genotypes of 52 inversions (45 generated by NAHR and 7 by NH methods) without tag SNPs in complete LD (*r*^2^ = 1) in the three studied super-populations (AFR, EUR and EAS). Inversion genotypes available in each population varied from 27 to 179 (with EAS having in most cases a lower number of samples), and MAF of polymorphic variants in a population ranged between 0.006 and 0.5, although 90% of the variants (47/52) show a MAF higher than 0.1 in at least one population (Supplementary Fig. 1). Inversion imputation accuracy was assessed by the *r*^2^ between the real and imputed genotypes in all the available samples where the inversions have been experimentally genotyped using the LOO strategy (see Materials and Methods). Specifically, we tested the performance of inversion phasing, five common imputation programs, filtering of imputed genotypes, or using a population-specific or global reference for imputation.

### Phasing

Since variant phasing is required for three of the imputation programs (Beagle, IMPUTE5, and Minimac4), and recommended for IMPUTE2, first we assessed whether imputation results differ between phasing methods. We found significantly higher imputation accuracy across inversions and populations when inversion phasing was done with Beagle compared to SHAPEIT5 in the four imputation programs tested (Supplementary Fig. 2). Therefore, only Beagle was used for phasing in all subsequent analyses.

### Imputation programs

Next, we tested inversion imputation performance of five imputation programs: Beagle Impute2 Impute5, Minimac4 and scoreInvHap (18–22). When considering all the inversions and populations together, the imputation accuracy varied considerably between methods. The best performing programs were IMPUTE5 and Minimac4, followed closely by IMPUTE2, while Beagle, and particularly scoreInvHap, show significantly worse results than the former two in all populations, except EAS (Fig. 1; Supplementary Table 4). These differences can also be appreciated in the distribution of the inversion imputation accuracy, with IMPUTE2, and, especially, IMPUTE5 and Minimac4 generating sharper peaks closer to *r*^2^ = 1 in most populations, whereas Beagle and scoreInvHap were able to impute perfectly a lower number of inversions (Fig. 1 and Supplementary Fig. 3).

**Figure 1.**
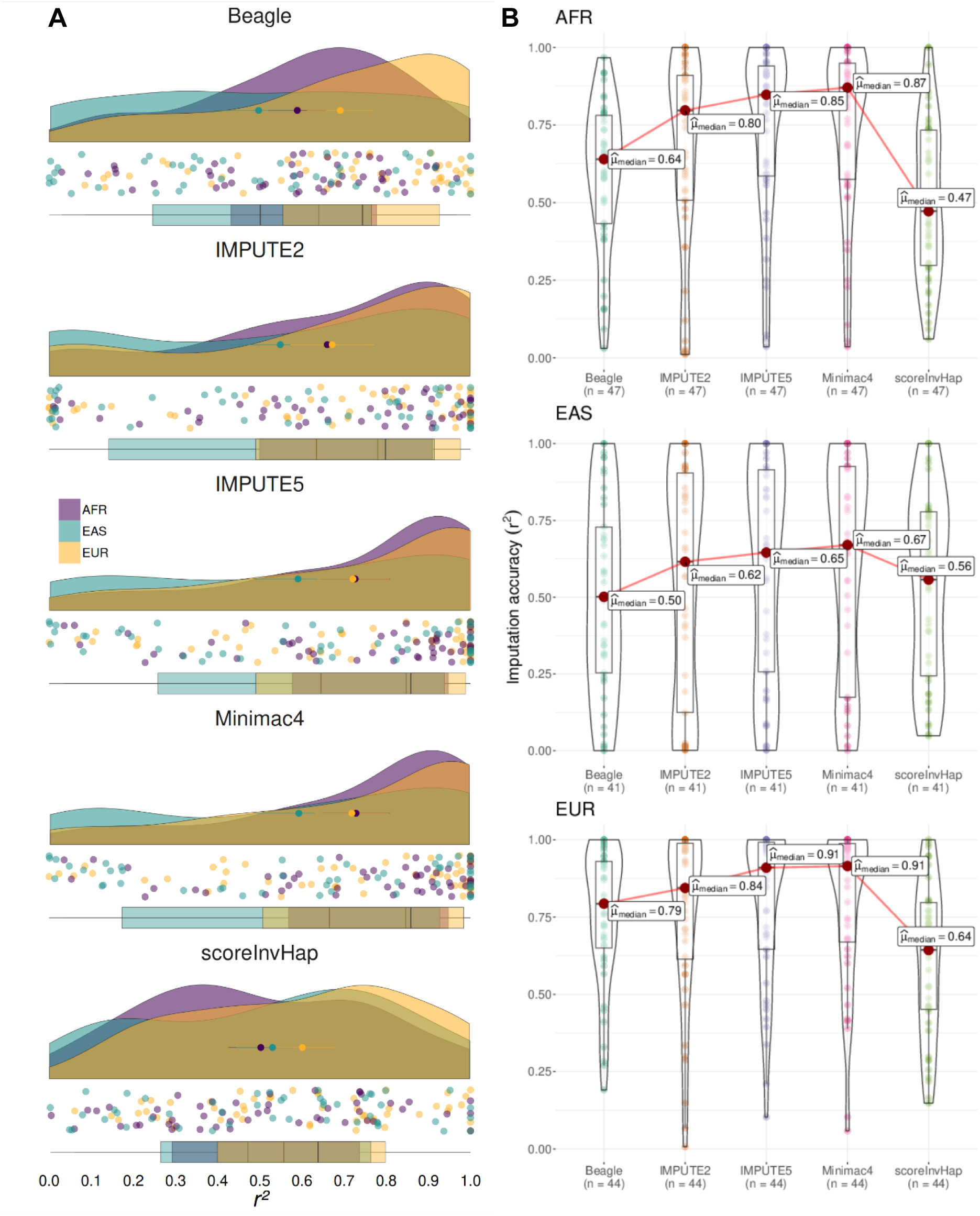
Inversion imputation results using different programs. (**A**) Distribution of original imputation accuracy values (*r*^2^) for the 52 analyzed inversions, divided by imputation program. Density plots in the three super-populations are represented together in different colors (purple for AFR, blue for EAS and orange for EUR), with the actual values for all inversions at the bottom (colored dots) and the mean and standard deviation in each population as dots with error bars within the graph. (**B**) Violin plots comparing the distribution of the imputation accuracy (*r*^2^) across the different programs in the three populations. Box plots represent the median (red dot) and interquantile range, with the values for all inversions indicated as colored dots. In each population, only inversions in which *r*^2^ can be calculated for all programs (*n*) are included in the comparison.

The distribution of inversion imputation accuracy also differed between populations, although, except for Beagle, no significant differences were found (Table 1 and Fig. 1). In general, inversions were imputed more accurately in EUR, followed by AFR. In contrast, EAS showed a wider *r*^2^ range that was biased to lower values for all programs (Supplementary Fig. 3), which is probably due to the reduced number of available genotyped samples in this case (Supplementary Fig. 1).

**Table 1.** Summary of inversion imputation results for different programs in three populations. For each method and population, the median imputation accuracy (*r*^2^) of the distribution of all the inversions that could be analyzed (*n*) is shown. Inversion imputation between populations was compared using an unpaired Wilcoxon rank-sum test and significant differences without multiple comparison correction are shown in boldface.

| Program | AFR | | EUR | | EAS | | Willcoxon rank-sum test $P$ value | | |
| --- | --- | --- | --- | --- | --- | --- | --- | --- | --- |
| | $n$ | $r^2$ | $n$ | $r^2$ | $n$ | $r^2$ | AFR-EUR | AFR-EAS | EUR-EAS |
| **Beagle** | 49 | 0.640 | 50 | 0.744 | 47 | 0.501 | **0.018** | 0.164 | **0.004** |
| **IMPUTE2** | 48 | 0.780 | 51 | 0.798 | 42 | 0.635 | 0.468 | 0.22 | 0.121 |
| **IMPUTE5** | 49 | 0.848 | 52 | 0.858 | 47 | 0.645 | 0.656 | 0.141 | 0.086 |
| **Minimac4** | 49 | 0.846 | 52 | 0.858 | 47 | 0.665 | 0.668 | 0.129 | 0.098 |
| **scoreInvHap** | 49 | 0.472 | 51 | 0.638 | 47 | 0.557 | 0.56 | 0.076 | 0.284 |

### Genotype probability filtering

As already mentioned, the GP is a score generated by each software that estimates the degree of confidence of the imputed genotypes. Except for scoreInvHap, the GP for every method tended to be high, with 83.7% (IMPUTE2) to 96.0% (Minimac4) of values exceeding 0.9 (Supplementary Fig. 4A).

To check the effect of GP filtering on inversion imputation, we screened a range of GP thresholds from 0.5 to 0.95 in the different programs (except for scoreInvHap, for which no filtering was applied due to its peculiar GP distribution). When considering the common set of inversions that can be analyzed by all programs, Minimac4 and IMPUTE5 show consistently the highest median imputation accuracy, with *r*^2^ values improving slightly with increasing GP (Fig. 2A). This effect is more drastic for IMPUTE2, with median imputation accuracy in the three populations reaching values closer or even higher (*e.g.* EAS) to the other two programs as GP threshold increases (Fig. 2A). However, IMPUTE2 improvement comes at a cost of excluding 8.4-17.4% low-confidence genotypes in the three populations at GP ≥ 0.90, comprising most samples for some inversions, which reduces the number of inversions with reliable results (Fig. 2A). Conversely, despite having high GP values similar to the other programs, median imputation accuracy in Beagle remained lower than in other methods across all GP filter thresholds (Fig. 2A).

**Figure 2.**
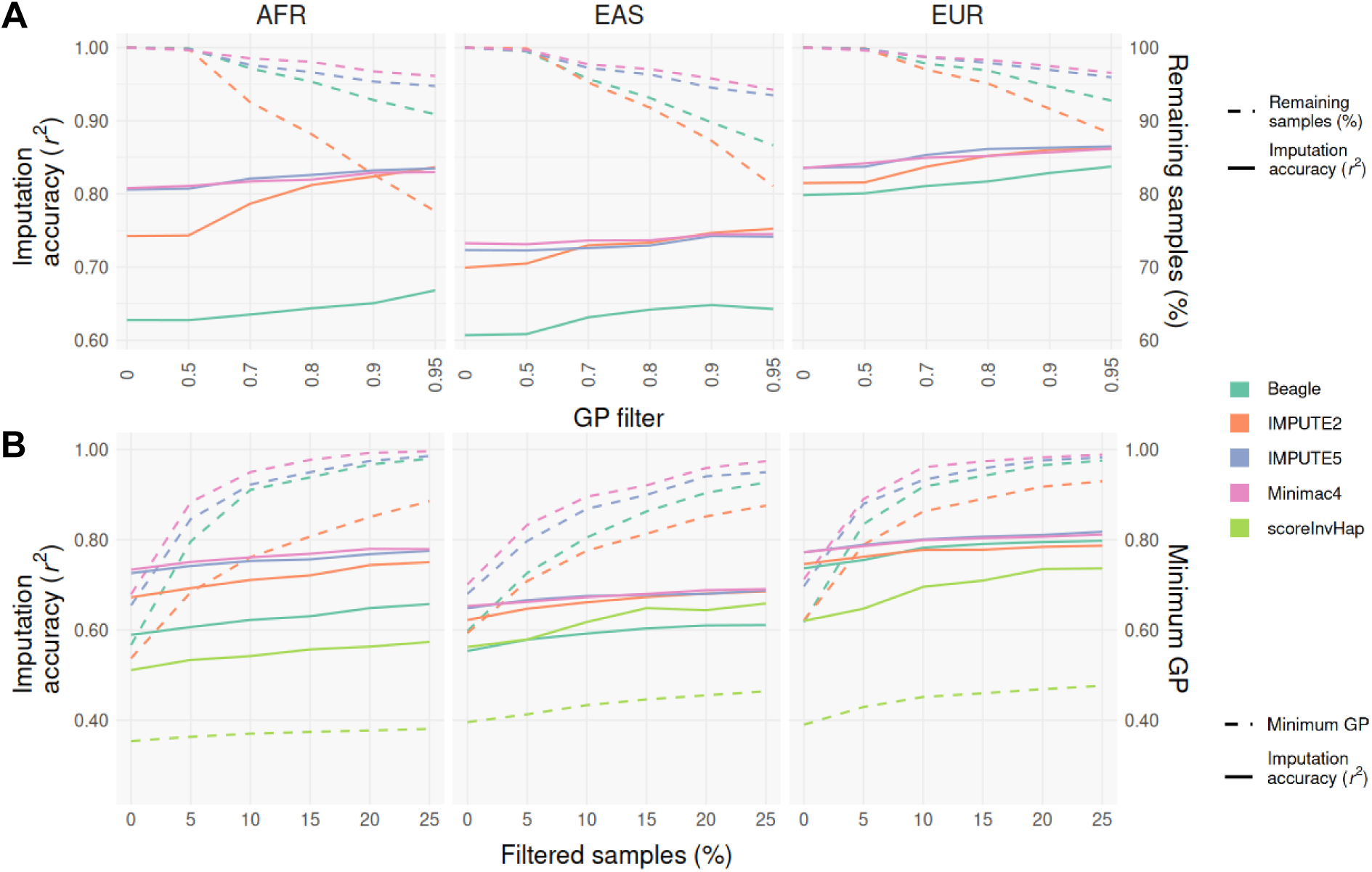
Effect of genotype probability (GP) filtering on inversion imputation accuracy (*r*^2^) with different imputation programs. (**A**) Median *r*^2^ (solid line) and sample loss (dashed line) in each population across increasing GP filtering criteria for all inversions together. (**B**) Median *r*^2^ (solid line) and lowest GP value of the remaining genotypes (dashed line) in each population for all inversions together, after removing progressively the worst imputed genotypes of every inversion (measured by the lowest GP) in 5% intervals (up to 25%). To make the results comparable, within each analysis and population we included only inversions that retain samples with both orientations at the strictest filtering criteria (GP ≥ 0.95 or 25% filtered samples), in which *r*^2^ with experimental genotypes can be calculated, across all the programs. In addition, those with less than 50% of samples remaining at the highest GP filter were excluded from the analysis. The number of analyzed inversions per population are: **A**, AFR = 38; EAS = 29; EUR = 38; and **B**: AFR = 44; EAS = 34; EUR = 43.

As already seen, GP values of the imputed genotypes varied among programs. Therefore, we also assessed the effect of filtering a fixed fraction of genotypes. For each inversion, 0% to 25% of the samples with the lowest GP were removed in increments of 5% and those inversions retaining both orientations at the strictest GP filter across all programs in each population were included in the comparison (Fig. 2B). Minimac4 and IMPUTE5 were again the best performing inversion imputation software for all filters, whereas IMPUTE2 showed lower median *r*^2^ in AFR and EUR when the same amount of samples were eliminated and similar values in EAS for ≥15% of filtered genotypes. On the other hand, Beagle and scoreInvHap had the lowest median *r*^2^, although filtering equal number of samples makes Beagle results get closer to those of IMPUTE2 and even surpass it in EUR. Similarly, sample filtering improved considerably scoreInvHap results, especially in EAS, in which it shows higher imputation accuracy than Beagle (Fig. 2B).

### Population versus global reference panel

Since some inversions were originally genotyped in a small number of samples of certain populations, like EAS, we also tested the effect on inversion imputation of using a global reference panel of all the samples of the three populations together (GLB), instead of just those from the population of the individual. Using a GP filtering of 0.9, the expansion of the reference size by combining different populations had overall a small effect on the results of the four programs that rely in a reference panel rather than in LD with SNPs (scoreInvHap), although it improved inversion imputability in certain situations (Supplementary Table 2). When looking at the distribution of the *r*^2^ difference between the GLB and population-specific reference, Beagle displayed considerable variation, followed by IMPUTE2, in which several inversions show clearly better imputation accuracy with the GLB panel, especially in EUR (Fig. 3; Supplementary Fig. 5). In contrast, for IMPUTE5 and Minimac4, a large proportion of inversions are imputed equally well in the GLB compared to the AFR or EUR population panels, although imputation accuracy with the GLB reference increases significantly in EAS (Fig. 3; Supplementary Fig. 5). In fact, as expected, inversions with higher imputation improvement in the GLB panel tend to be those with lower population sample size (Supplementary Table 2).

**Figure 3.**
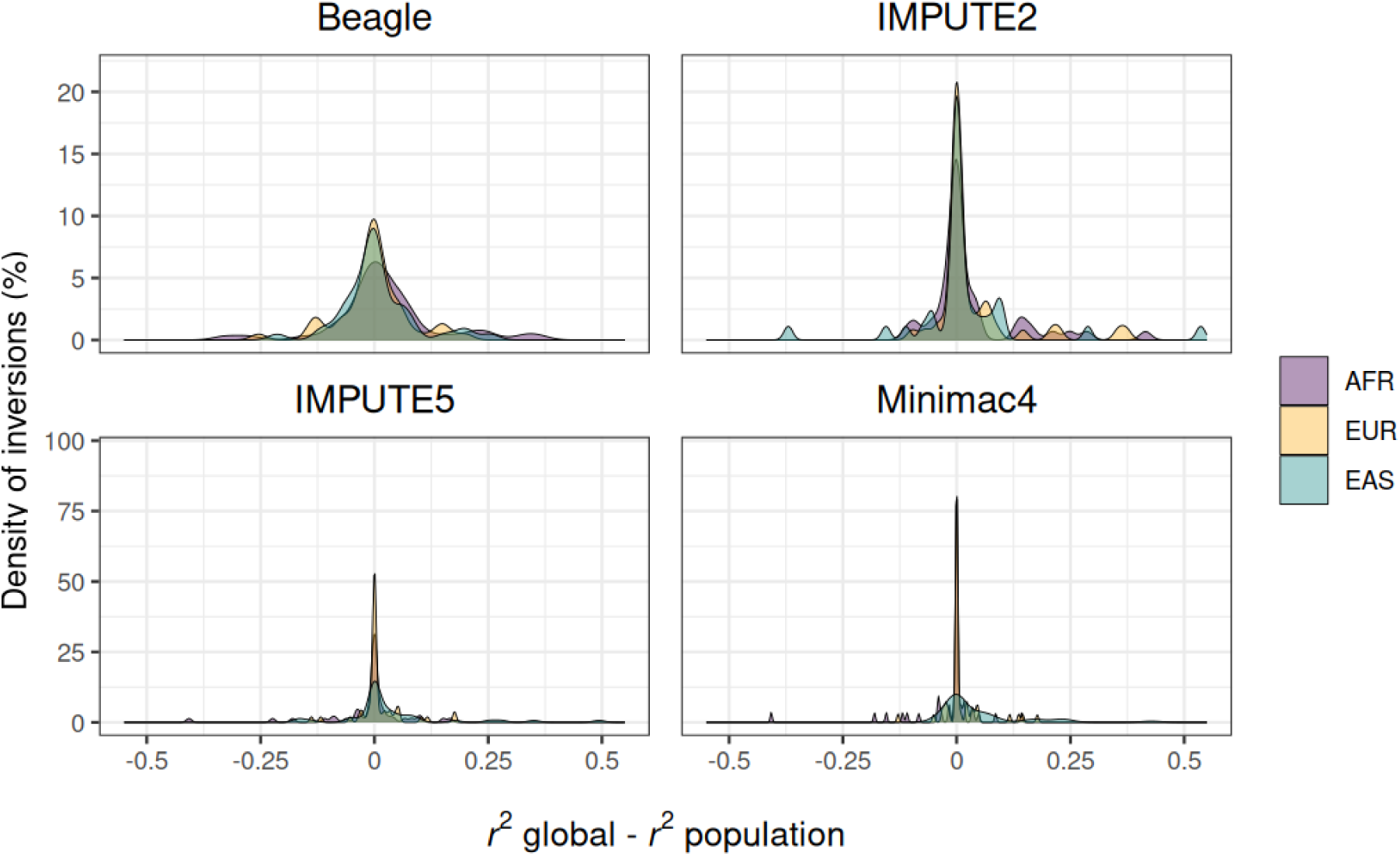
Comparison of imputation accuracy (*r*^2^) between using different reference panels. The graphs show the density distribution (Y axis) of the difference in *r*^2^ of the global minus the population panel across all analyzed inversions (X axis) for four imputation programs, with the three populations represented in different colors. Individual graphs for each population and additional data are shown in Supplementary Fig. 5.

### Inversion imputation results

Next, we assessed how many of the analyzed inversions could be imputed accurately from WGS data in each population (defined as those with good imputation accuracy (*r*² ≥ 0.8) and more than 50% of samples remaining after filtering). Using the population-specific reference panel, excluding low-confidence genotypes in general increased the number of imputable inversions, with the GP ≥ 0.9 filter typically outperforming the no filter and GP ≥ 0.8 filter results in most programs (Fig. 4A). However, in Minimac4 and IMPUTE2 the distribution of GP values is skewed towards higher values (Supplementary Fig. 4) and the number of imputable inversions tends to be very similar across GP filters. Conversely, scoreInvHap accurately imputes very few inversions without filters (12.8-23.1%) and GP filtering removes virtually all the imputed genotypes (Fig. 4A). Therefore, except for scoreInvHap, a GP ≥ 0.9 filter was established as the standard criteria for inversion imputation.

**Figure 4.**
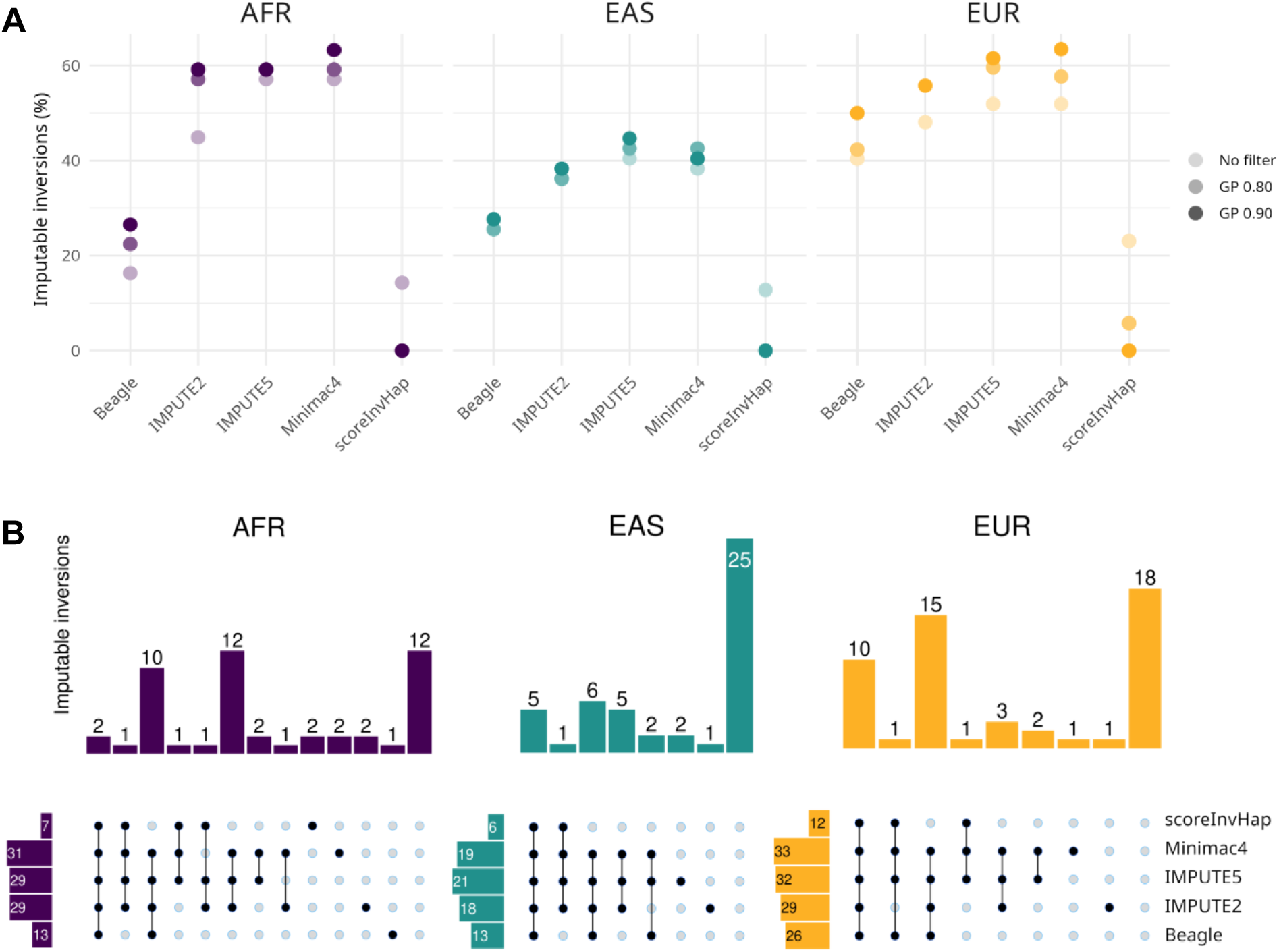
Inversion imputability with different programs in three populations. (**A**) Effect of GP filtering (including no filter and GP ≥ 0.8 or 0.9) on the number of inversions that can be imputed by each program (*r*^2^ ≥ 0.8 and ≥50% remaining samples), expressed as the percentage of the inversions having more than two chromosomes with the minor orientation in the analyzed samples of that population. (**B**) UpSet plots showing the intersection of imputable inversions across different imputation tools after applying a GP ≥ 0.9 filter (except for scoreInvHap) in the three populations. Each column represents a unique combination of programs that successfully imputes a given inversion and the number of imputed inversions in that combination (bar chart). Left bars indicate the total amount of inversions imputed by every method.

Considering the 0.9 GP filter, imputable inversions in each population with the four best performing programs ranged from 27.7% in EAS (13) for Beagle to 63.5% in EUR (33) for Minimac4. Minimac4 and IMPUTE5 were the programs imputing the highest number of inversions in EUR and AFR or in EAS, respectively, with IMPUTE2 being close behind (Fig. 4A). Beagle, on the other hand, could only impute reliably 27.7-50% of the inversions. When combining the results of all the programs, the total number of imputable inversions increased to 75.5% in AFR (37) and 65.4% in EUR (34), while consistent with the lower imputation accuracy (Fig. 1; Fig. 2), only 46.8% of inversions (22) could be accurately imputed in EAS (Fig. 4B). Looking into the imputability at inversion level, a good consistency was found between methods, and most inversions could be accurately imputed by at least two of them, with Minimac4 and IMPUTE5 being the biggest contributors and imputing basically the same inversions (Fig. 4B). Nevertheless, there was a population bias in the distribution of tools imputing an inversion. Whereas in EAS and EUR the vast majority of imputable inversions (86.4% and 88.2%, respectively) could be imputed by more than two methods (and many by even four or five), in AFR there is more variation between programs and this number reaches only 73.0% (Fig. 4B), probably due to the higher haplotype diversity that makes imputation more challenging.

Individual results of the different imputation methods are shown in Supplementary Table 2 and Supplementary Fig. 6. In general, the majority of programs obtained comparable imputation accuracies for each inversion and population, with small variations resulting in inversions being imputable in some cases but not in others. The average Pearson correlation coefficient (*R*) between the *r*^2^ values of Beagle, IMPUTE2, IMPUTE5 and Minimac4 compared to the other four methods is, respectively, 0.73, 0.74, 0.78 and 0.77, with the last two being the most similar (*R* = 0.98). The main exception was again scoreInvHap, which tends to generate the most different imputation results with respect to the rest of methods (average *R* between *r*^2^ values of 0.42) (Supplementary Fig. 6). When considering the best method for the imputable inversions, again three tools clearly stand out, with more inversions being imputed more reliably by IMPUTE2 in AFR, IMPUTE5 and Minimac4 in EAS, and Minimac4 in EUR (Supplementary Table 5), although differences in imputation accuracy tend to be small. Specifically, in each population, between 22.4% and 31.9% of the inversions were imputed with *r*^2^ = 1 by at least one method (11 in AFR, 15 in EAS and 16 in EUR), with IMPUTE2, IMPUTE5 or Minimac4 always imputing perfectly the highest number of inversions (Supplementary Table 5). However, IMPUTE2 results always come at the cost of retaining less samples (Supplementary Table 2). If a global reference panel is used instead, similar patterns are observed and improvements in imputation accuracy tend to be minor, but there are three extra inversions showing *r*^2^ = 1 imputation accuracy in certain populations, including two in EAS that could not be imputed accurately before. Interestingly, two inversions (HsInv0290 and HsInv1111) could be imputed well with scoreInvHap in AFR, while very bad results were obtained for them with the other methods (Supplementary Fig. 6). These inversions have in common that they span a relatively large region including big IRs, although scoreInvHap performance varies among populations.

### Factors affecting imputation accuracy

Besides the different programs, several factors can affect the imputation of individual inversions. For example, all NH inversions, which have a unique origin, could be accurately imputed (usually with *r*^2^ = 1) by most programs in the analyzed populations using the local or global reference panel (Supplementary Table 2 and Supplementary Fig. 6). On the other hand, NAHR inversions, which tend to be recurrent, show worse imputation results (Supplementary Table 2; Supplementary Fig. 6), with 46.7% (21/45) not accurately imputed in any or just one of the studied populations. However, another 31.1% (14/45) can be imputed well in all the populations in which the inversion is present with at least one program, although imputation accuracies are much lower than for NH inversions (Supplementary Table 2 and Supplementary Fig. 6). Actually, NAHR inversion imputation *r*^2^ values for IMPUTE5 and Minimac4 are positively correlated with the maximum LD with other variants in each population, which is a good approximation of how many chromosomes are affected by recurrence events (IMPUTE5: *R* = 0.504, *N* = 124, *P* = 2.46 x 10^-9^; Minimac4: *R* = 0.502, *N* = 125, *P* = 2.49 x 10^-9^). In addition, some NAHR inversions are imputable in certain populations in which there are no or few recurrent chromosomes, but not in others with higher levels of recurrence (e.g. HsInv0390, HsInv0395 and HsInv0605 in EUR vs. AFR) (Supplementary Fig. 6). Finally, another factor with a significant effect on imputability is variant frequency, especially for recurrent inversions, with more variability and worse imputation accuracy in populations with low versus high frequency (e.g. HsInv0341 in AFR vs. EUR and EAS). Nevertheless, imputation results likely depend much on other characteristics related to the genetic structure of the region, with several poorly imputed inversions across populations despite showing relatively high LD, such as HsInv0608, which has one of the highest recurrence rates (11). Conversely, there are other inversions showing good imputation accuracy despite having lower LD levels, like HsInv0390 in EUR or HsInv0114, HsInv0266, HsInv0396 and HsInv0403 in AFR.

To better quantify the influence of several biological features on imputability of NAHR inversions, we fitted linear models of each program imputation results (except scoreInvHap) using as variables the inversion MAF, number of SNPs in the region, type of chromosome (Chr. X vs autosomes), number of genotyped samples, and ratio between the IR and the inversion size (IR/inversion length), which has been previously associated with inversion recurrence (10, 11). Without GP filtering, which allows us to compare basically the same inversions, the different programs gave similar results, with variables explaining between 12.8% and 22.5% of the variance in imputation accuracy values (Supplementary Table 3). Removing imputed genotypes with GP < 0.9 improved slightly the variance in imputation accuracy explained by most models, which almost doubled for IMPUTE2 (Supplementary Table 3), probably due to removing some of the most complex inversions. Apart from this, the only variable that had a consistent significant effect is the number of genotyped samples, increasing imputation accuracy in all cases (Fig. 5). Other variables that have been associated to higher inversion recurrence levels (10, 11) showed consistent effects across models, but they were only significant in a few of them. For example, in general, IR/Inversion length ratio and Chr. X inversions were negatively correlated with imputation accuracy, although results were only significant for IMPUTE2 and, to a lesser extent, IMPUTE5 in some of the models (Fig. 5; Supplementary Table 3). Finally, MAF and the number of SNPs tend to have small opposite effects on imputation, with significant negative results for the latter with GP filtering for IMPUTE2.

**Figure 5.**
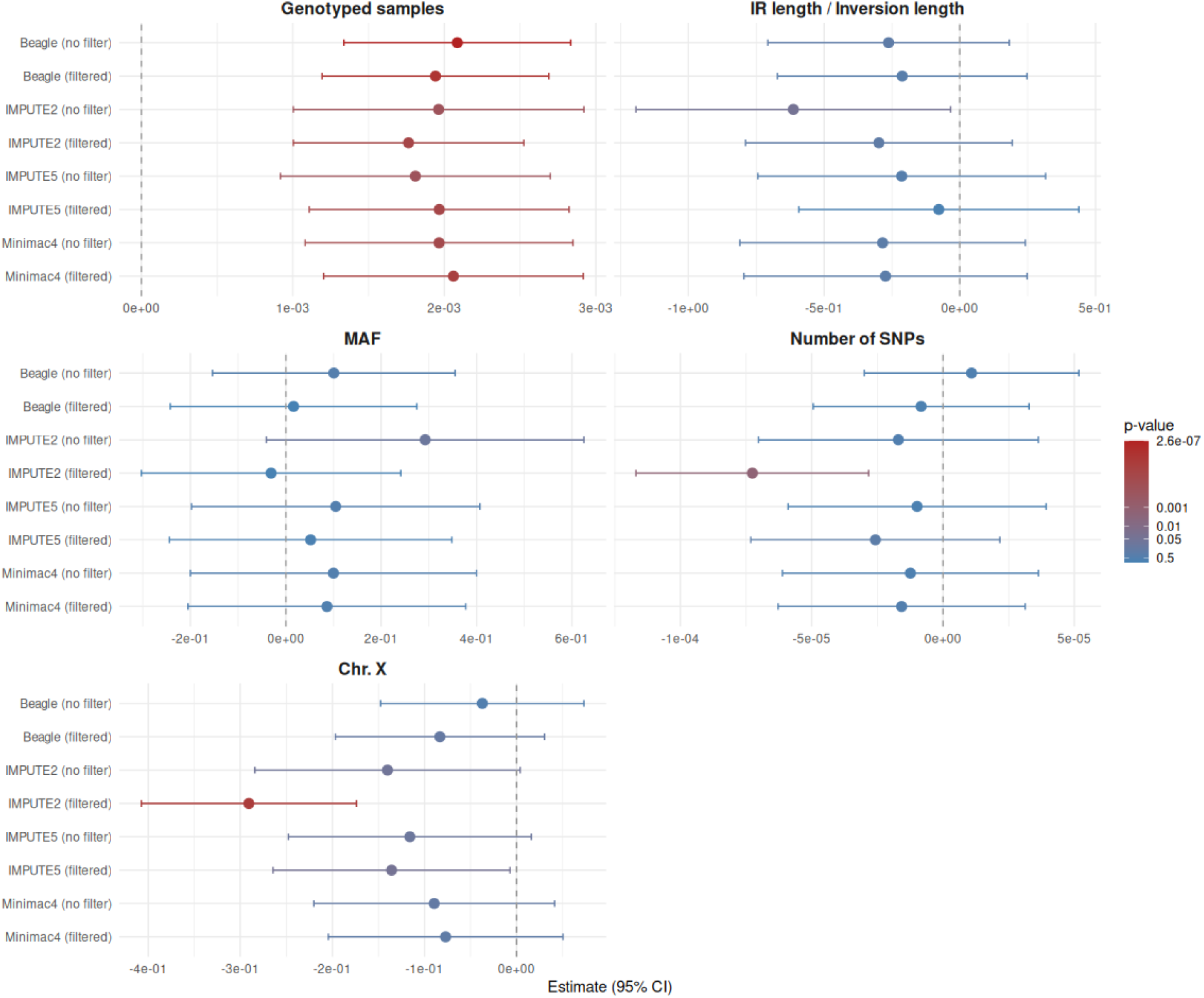
Results of linear models of NAHR inversion imputation using different programs. Graphs show the model coefficients (X axis) of the different variables in each program model (Y axis), with and without removing unreliable imputed genotypes having genotype probability (GP) values lower than 0.9 (labeled as filtered and no filter, respectively). Error bars represent the 95% confidence interval (CI) and color indicates the significance of the variable effects according to the p-value scale (right side). Dashed line marks the 0 value (no effect).

### Imputation from SNP array data

As the last step, we checked how well inversions are imputed with SNP genotyping arrays and compared the results with those from WGS. As mentioned in the introduction, arrays interrogate fewer variants, which could significantly affect imputation. To test this, we simulated two types of commonly used SNP arrays with different density: Omni high-resolution array (∼2.4 million SNPs) and GSA low-resolution array (∼650,000 SNPs). In general, reliability of the genotypes obtained with the different imputation programs was proportional to the number of SNPs available, which reduces considerably the total fraction of imputed inversion genotypes with GP ≥ 0.9 in the different programs to 77.5-85.8% for the Omni and 66.6-78.6% for the GSA arrays (Supplementary Fig. 4). As there are less variants in the reference panel, the total number of imputable inversions in the three populations also decreased to 42.6-65.3% for Omni and 40.4-59.5% for GSA (Fig. 6). In addition, median imputation accuracy with most programs was clearly lower for GSA than Omni or WGS data in EAS and AFR, although in EUR the differences were smaller (Fig. 6A). However, Omni and WGS tend to behave more similarly, with median imputation accuracy being equal or even higher for Omni in some populations and programs (Fig. 6A), suggesting that there is not so much benefit of a greater density of SNPs. The main exception was Beagle, which, surprisingly, in the three populations showed higher imputation accuracy and number of imputed inversions with both arrays than with WGS data (Fig. 6).

**Figure 6.**
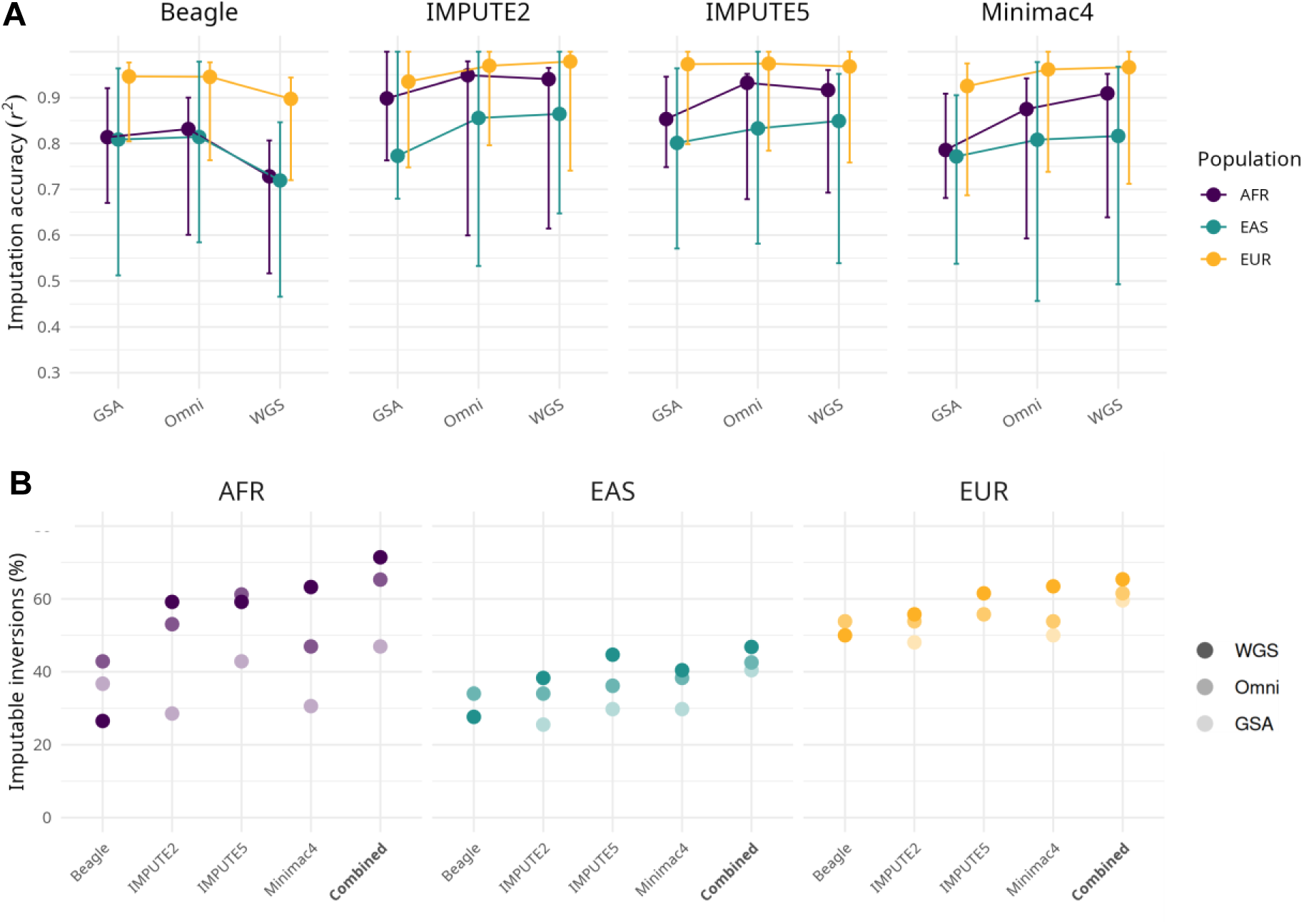
Inversion imputation results with SNP array data. (**A**) Median imputation accuracy (*r*^2^) across inversions from Omni and GSA array data compared to whole genome sequencing (WGS) using four imputation programs after GP ≥ 0.9 filtering in the three analyzed populations. To make the results comparable, within each population we included only inversions that retain samples with both orientations in all the analyses: AFR = 35; EAS = 28; EUR = 37. (**B**) Number of inversions that can be imputed by each program (*r*^2^ ≥ 0.8 and ≥50% remaining samples) with the different types of array and WGS data, expressed as the percentage of the inversions having more than two chromosomes with the minor orientation in the analyzed samples of that population. The total number of inversions imputable by any of the four programs is shown in the Combined category.

When considering the different populations, results were somewhat variable between the tested arrays and programs (excluding Beagle). As before, EUR showed the highest imputation accuracy and there was little variation between the different arrays and WGS, with high inversion imputability rates even in the smaller array (Fig. 6). In EAS, imputation results from the three types of data were also relatively similar, although they were much lower than in the other populations, as happens for WGS data (Fig. 6). Conversely, in AFR imputation accuracy and the number of imputable inversions were more or less equivalent between WGS and the Omni array, excluding Minimac4, but GSA exhibited a quite lower number of imputable inversions (Fig. 6), which suggests that the array SNPs do not properly capture African genetic diversity. Despite this variation, overall IMPUTE5 was the best performing method with the array data, with IMPUTE2 close behind, whereas Minimac4 appeared to be more hindered by the SNP sparsity, especially in EUR and AFR.

## Discussion

Accurate genotypes are crucial to determine the functional and phenotypic effects of genetic variants. By performing for the first time an exhaustive comparison of the imputation of challenging inversions using different methods, we found that 46.8-75.5% of the studied inversions in each population can be imputed with high accuracy from WGS data, despite our relatively strict (*r*^2^ ≥ 0.8) imputability criteria. Therefore, this work paves the way to generate reliable information for a higher fraction of inversions that goes beyond previous analyses based on a single imputation tool (10, 23, 28). Nevertheless, there are still many inversions remaining that cannot be imputed well with any program in either all or certain populations. In addition, we have found differences in imputation accuracy between populations, with worse imputation in AFR and especially EAS, which suggest that imputation success relies heavily on the haplotype diversity of the population in the affected regions and the number of available reference genotypes to properly capture this diversity.

Imputation algorithms differ in their approaches to modeling haplotype inheritance (18–22). Inversions represent a special type of variant since they suppress recombination (9), which may disrupt these calculations, affecting their own imputation. Here, we have shown that IMPUTE5 and Minimac4 outperformed the other tools, with IMPUTE5 performing slightly better in the array data, making them our recommended choices for inversion imputation. Although IMPUTE2 achieves comparable imputation results for many of the inversions (Fig. 4; Supplementary Table 3), it does so at a significant cost in sample loss following GP filtering. This effect is particularly pronounced for low MAF inversions (Supplementary Table 2). Moreover, IMPUTE2 has much longer running times and requires more computing resources than the other tools, limiting its utility. In contrast, the poor results of scoreInvHap may stem from its direct reliance on LD between inversions and surrounding variants, which likely undermines its performance in this dataset. Nevertheless, there are a few large inversions that are accurately imputed only with scoreInvHap, as seen before for the 4.2 Mb 8p23.1 inversion (29). This suggests that for large inversions the other methods may be negatively affected by positioning the inversion in the reference panel as its middle point, since they take into account the local variation around the variant, which could be quite distant from the actual breakpoints. Besides, there is no clear explanation for the relatively low inversion imputation of Beagle, although it could be related to the higher DNA sequence fragmentation of this method, which could compromise the imputation of larger SVs compared to SNPs. In any case, as shown for scoreInvHap, there is no single best imputation method for all inversions, and it might be necessary to use several ones to impute as accurately as possible the maximum number of variants.

Aside from testing the different programs, we also assessed several other alternatives that can affect imputation. In particular, GP filtering markedly improved imputation accuracy, while minimizing sample loss (Fig. 2). Since imputation is typically a preparatory step for functional analyses, excessive sample filtering can reduce statistical power or introduce bias toward specific haplotypes. The top-performing methods, IMPUTE5 and Minimac4, exhibited acceptable sample retention, with only 5.9% and 4% of samples lost, respectively, in WGS data after a GP ≥0.9 filtering, which is much higher for IMPUTE2 (16.3%) (Supplementary Fig. 4A). Therefore, the 0.9 GP criteria seems a good compromise between imputation accuracy and sample size. On the other hand, using a global reference panel combining multiple populations could help imputation in some specific inversions with low sample size (such as in EAS), but gains compared to a population specific panel tend to be small and in some cases imputation accuracy can decrease due to combining recurrence events from different populations.

Among the biological factors examined, recurrence represents clearly the biggest challenge for imputation, since both inversion orientations can coexist in exactly or virtually identical haplotypes (10, 11). In fact, despite showing good imputation accuracies, genotypes filtered out due to low GP in some inversions are likely caused by recurrent events (such as in HsInv0193, HsInv1117 or HsInv0286). Apart from this, as seen already for EAS, the number of samples was the only variable significantly affecting imputation in all models (Fig. 5). Other factors had also some influence on imputation, although in most cases they were not significant (Fig. 5). For example, the number of SNPs tends to have a negative effect, which might be related to the size of the inversion and the distance between the imputed variable position and the actual inversion breakpoints, whereas MAF consistently had a positive effect, supporting the need of certain amount of haplotypes of both orientations for the programs to be able to detect and impute them correctly. In the case of chromosome type, Chr. X inversions exhibited lower imputation accuracy across all methods, but the difference does not appear to stem from disparities in imputation performance between autosomal and sex chromosomes (16). Instead, the explanation likely lies in NAHR-mediated inversion hotspots created by the elevated frequency of segmental duplications and the high inversion recurrence rate observed in Chr. X (10–12, 30). Nevertheless, the ratio between the size of the inversion and the IRs, which is a good predictor of recurrence levels (10, 11), showed only a moderate negative effect on imputation in most methods. This is probably due to the opposite effects of inversion length, both hampering imputation and decreasing recurrence, and other factors that could affect imputation accuracy, like the evolutionary history and the number of individuals affected by each recurrence event.

However, despite the inversion imputation improvements, there are some important limitations. First, for some inversions the number of available genotypes is quite limited, which reduces the accuracy of the imputation, especially in certain populations. More importantly, as already mentioned, a remarkable proportion of inversions are still not well imputed with any of the methods, mainly due to high recurrence levels. Resolving such complexity requires a larger reference panel to include enough representatives of all haplotype combinations (31). Current efforts to identify IR-mediated inversions using ultra-long reads are expected to generate more exhaustive genotype data for most human inversions, including many of those most challenging (8, 32–34), which will result in better imputation of a larger number of inversions. In addition, future methodological developments, such as the application of machine learning strategies, could also help impute recurrent events. Therefore, our results shed light on the capabilities and limitations of existing tools for handling structural variants, setting the stage for further exploration of imputation strategies in complex genomic contexts. Such advances should ultimately allow us to explore in detail the functional effects and phenotypic association of the whole set of human genomic variation.

## Data availability

All the data generated in this article are available in the article and in its online supplementary material. Inversion genotypes are available in dbVar (Study accession number nstd169 and nstd185) and 1000 Genomes High Coverage data can be obtained from https://ftp.1000genomes.ebi.ac.uk/vol1/ftp/data_collections/1000G_2504_high_coverage/working/20220422_3202_phased_SNV_INDEL_SV/.

## Supplementary data statement

Supplementary Data are available at NAR Genomics and Bioinformatics Online.

## Acknowledgements

We thank J. Lerga-Jaso for advice and help with imputation and data analysis, and the Comparative and Functional Genomics group members for the inversion experimental genotypes and valuable comments.

## Author contributions statement

MC conceived the project and supervised the different steps. IY and AM developed the methodology and performed the analyses. All the authors contributed to the interpretation of the results and the writing of the manuscript.

