## Supplementary Figures for "Accurate imputation of inversions in human genomes using different algorithms and data sources"

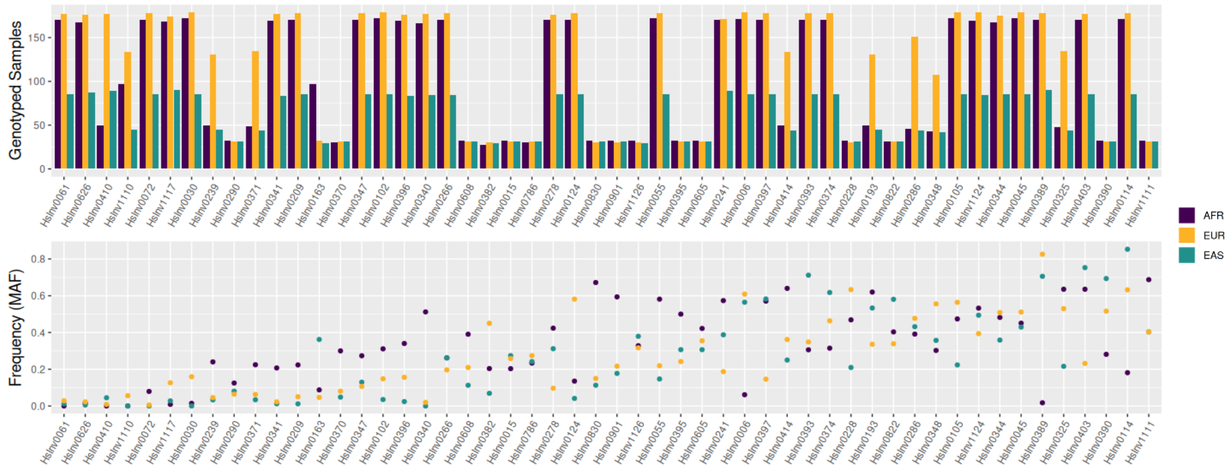


**Supplementary Figure 1.** Summary of available information for the 52 inversions analyzed, including the number of experimentally genotyped samples (top) and the minor allele frequency (MAF, bottom) in the African (AFR), European (EUR) and East-Asian super-populations.

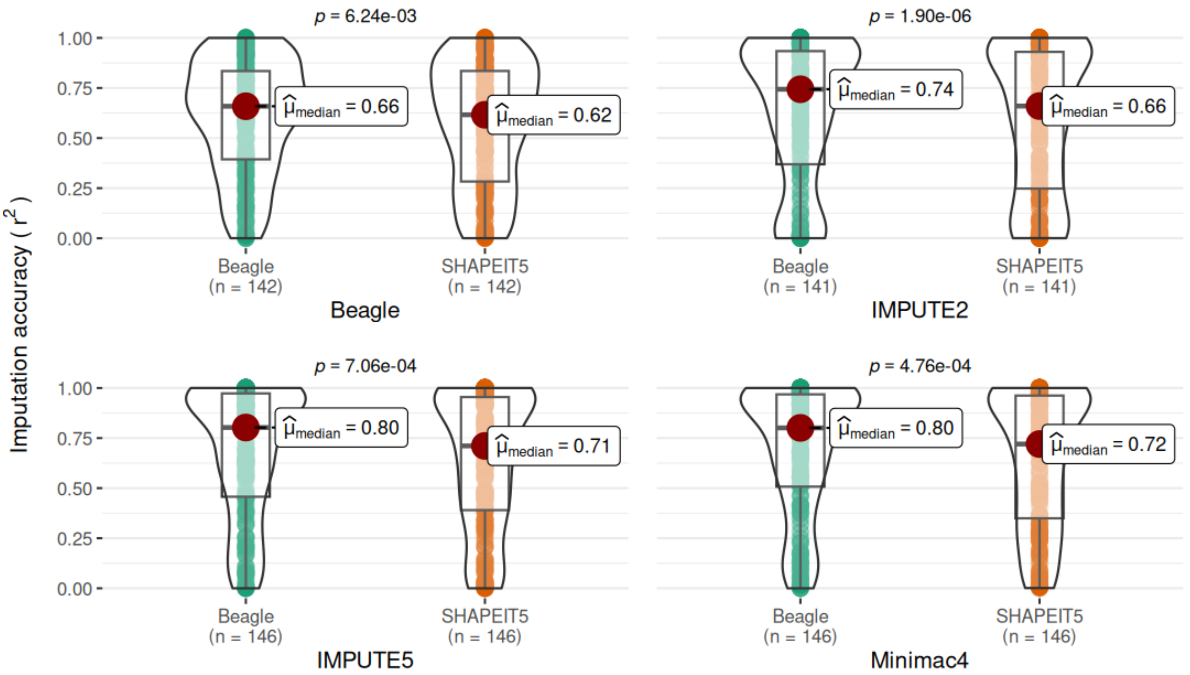


**Supplementary Figure 2.** Comparison of inversion imputation results with different phasing and imputation programs. Violin plots represent the imputation accuracy ($r^2$) of the 52 analyzed inversions in each of the three super-populations with four imputation programs, after inversion phasing using Beagle (green, left) or SHAPEIT5 (orange, right). Only those imputed inversions with variable orientations in a population in both phasing programs in order to be able to calculate the $r^2$ are included (indicated by $n$) and the median $r^2$ value across inversions and populations is shown inside the rectangle (μ). *P*-value ($p$) of the difference in imputation accuracy between Beagle and SHAPEIT5 is shown in top of each comparison and was calculated by a paired Wilcoxon signed-rank test.

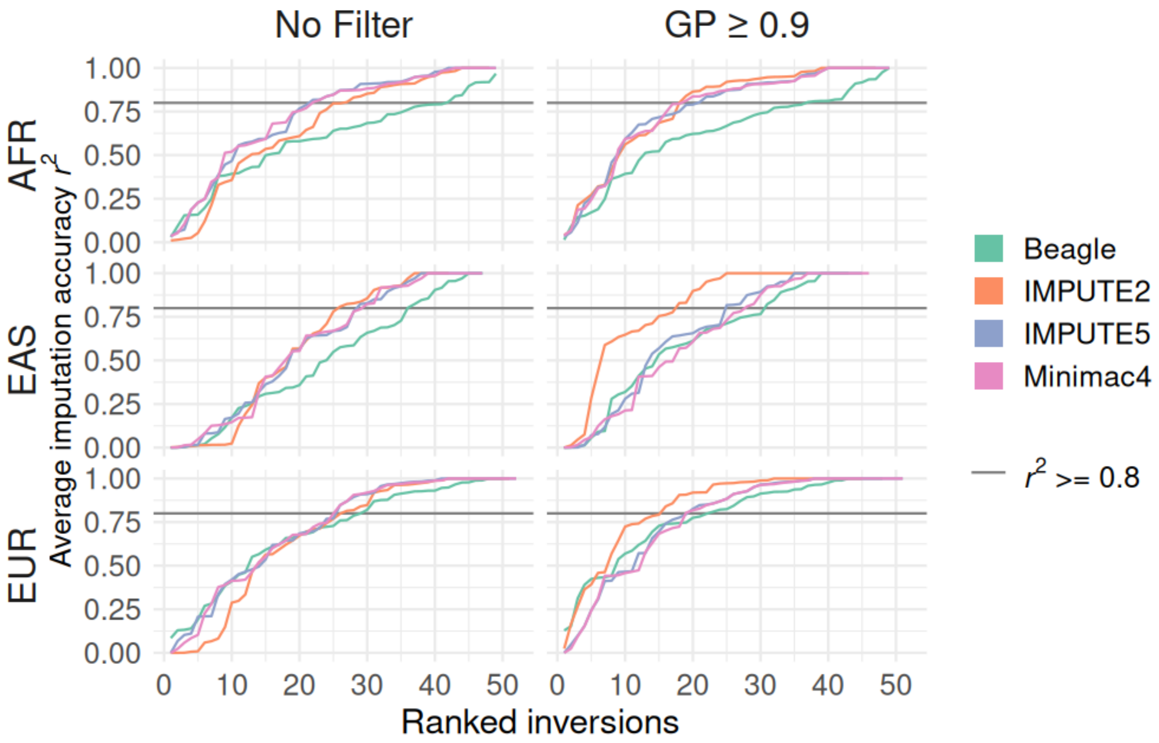


**Supplementary Figure 3.** Distribution of inversion imputation accuracy ($r^2$) for four of the programs tested. Inversions are ranked according to the $r^2$ across all samples of each population (y axis), from the lowest to the highest value per method (including inversions in which $r^2$ cannot be calculated due to lack of imputed minor alleles as 0). Both the original $r^2$ values (left) and those after filtering out imputed genotypes with probability (GP) lower than 0.9 (right) are represented. The faster the $r^2$ = 0.8 imputability threshold is reached (represented by a solid solid), indicates that there is a higher number of imputable inversions. Conversely, with Beagle a lower proportion of imputable inversions is obtained in all cases. IMPUTE2 shows better performance than other methods with the 0.9 GP filter, although this is based on less genotypes for many inversions and a smaller number of easier to impute inversions.

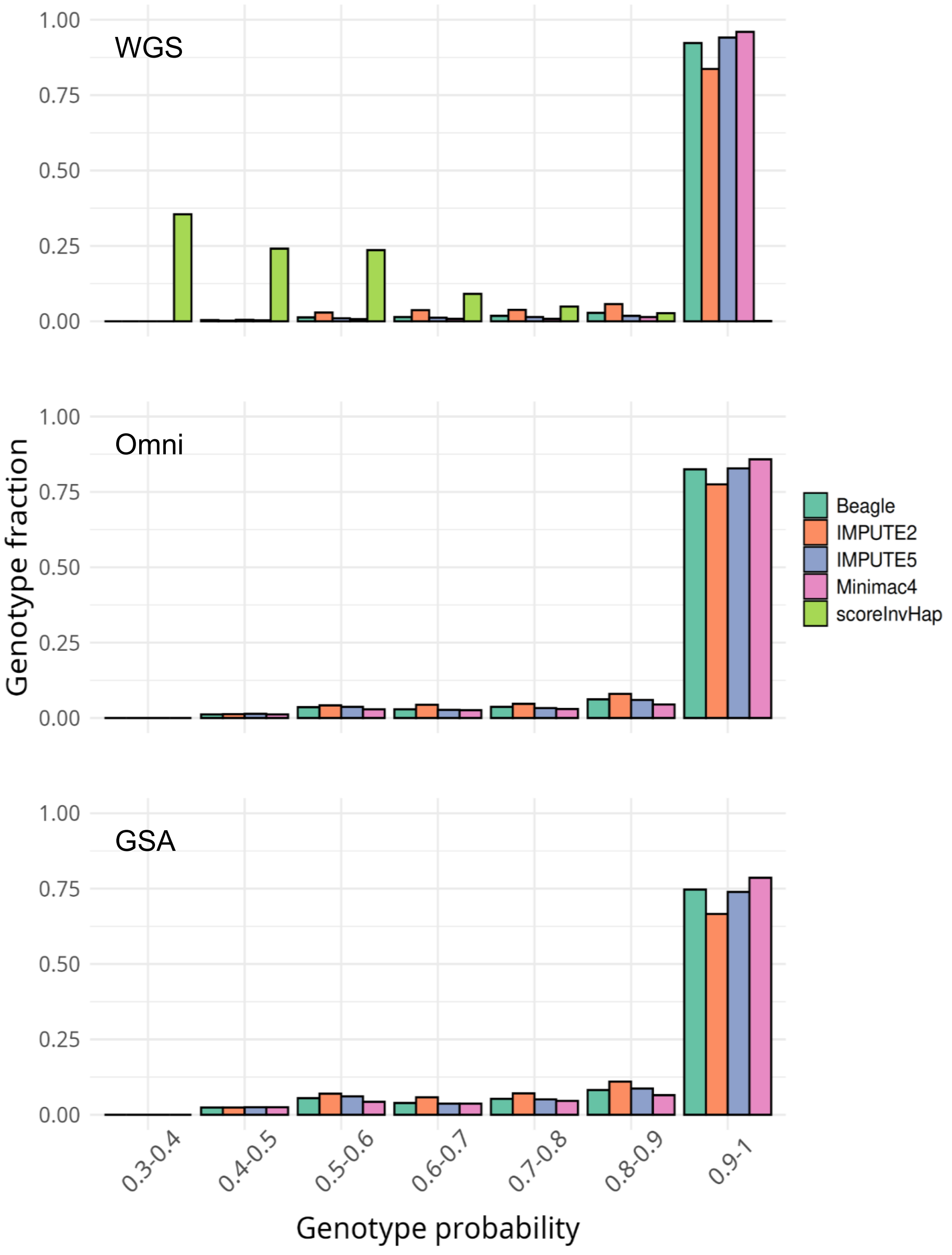


**Supplementary Figure 4.** Histogram of the distribution of genotype probability (GP) values of imputed genotypes obtained by each program using nucleotide variation data from whole genome sequence (WGS) or Omni and GSA SNP genotyping arrays. Fraction of genotypes is calculated in GP bins of 0.1 combining all samples from different populations and inversions together.

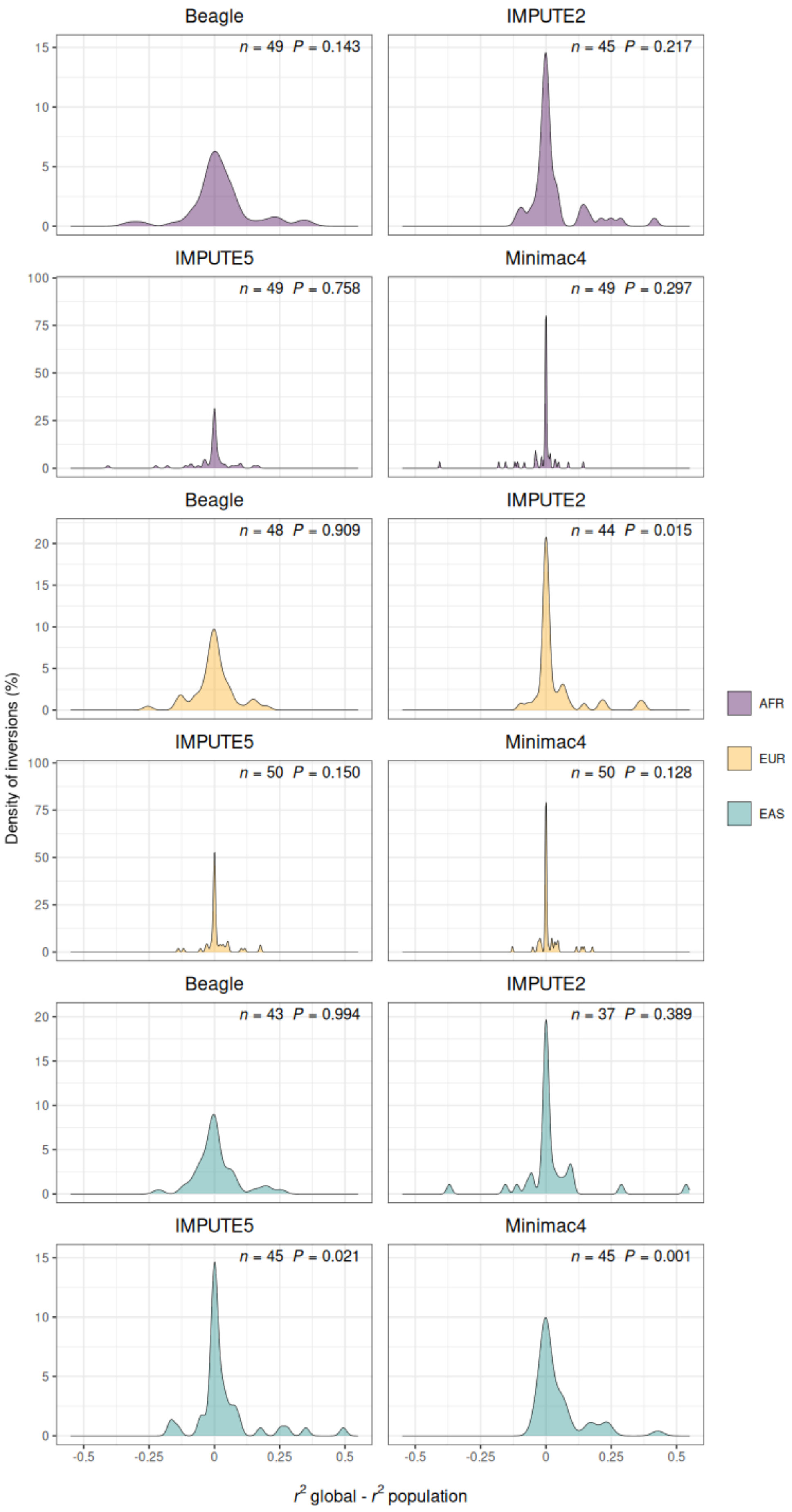
Beagle
n = 49 P = 0.143
IMPUTE2
n = 45 P = 0.217
IMPUTE5
n = 49 P = 0.758
Minimac4
n = 49 P = 0.297
Beagle
n = 48 P = 0.909
IMPUTE2
n = 44 P = 0.015
IMPUTE5
n = 50 P = 0.150
Minimac4
n = 50 P = 0.128
Beagle
n = 43 P = 0.994
IMPUTE2
n = 37 P = 0.389
IMPUTE5
n = 45 P = 0.021
Minimac4
n = 45 P = 0.001
Density of inversions (%)
r2 global - r2 population
AFR
EUR
EAS

**Supplementary Figure 5.** Detailed comparison of imputation accuracy ($r^2$) between using different reference panels with four imputation programs. Each graph shows the density distribution (Y axis) of the difference in $r^2$ of the global minus the population panel across all analyzed inversions (X axis) for one population and imputation program, with the three populations represented in different colors. The number of inversions (*n*) analyzed in each case and the *P* value of a Wilcoxon paired signed-rank test to compare the $r^2$ of the two reference panels are shown inside the graphs.

### Imputation method performance per inversion in AFR population

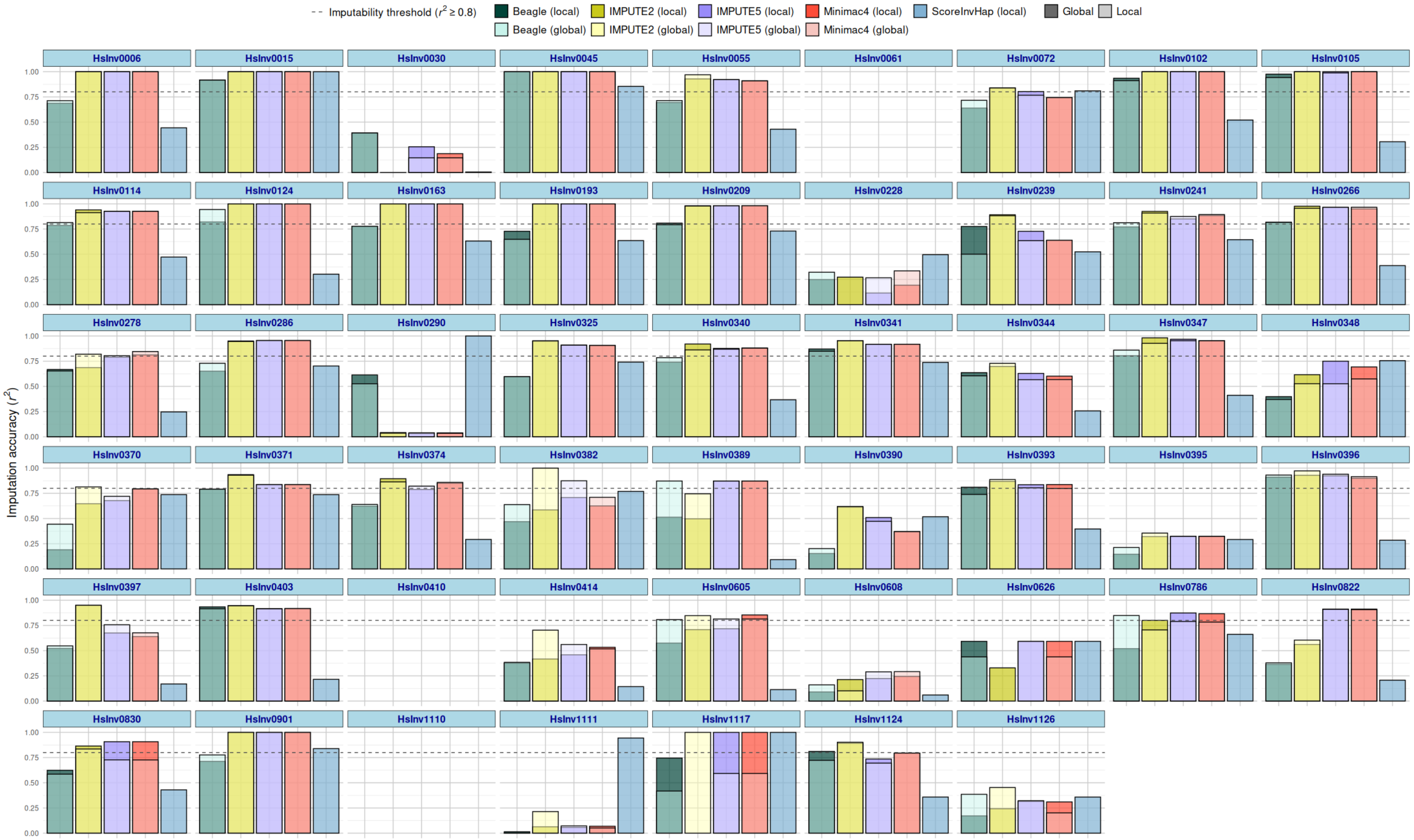

**B**

### Imputation method performance per inversion in EUR population

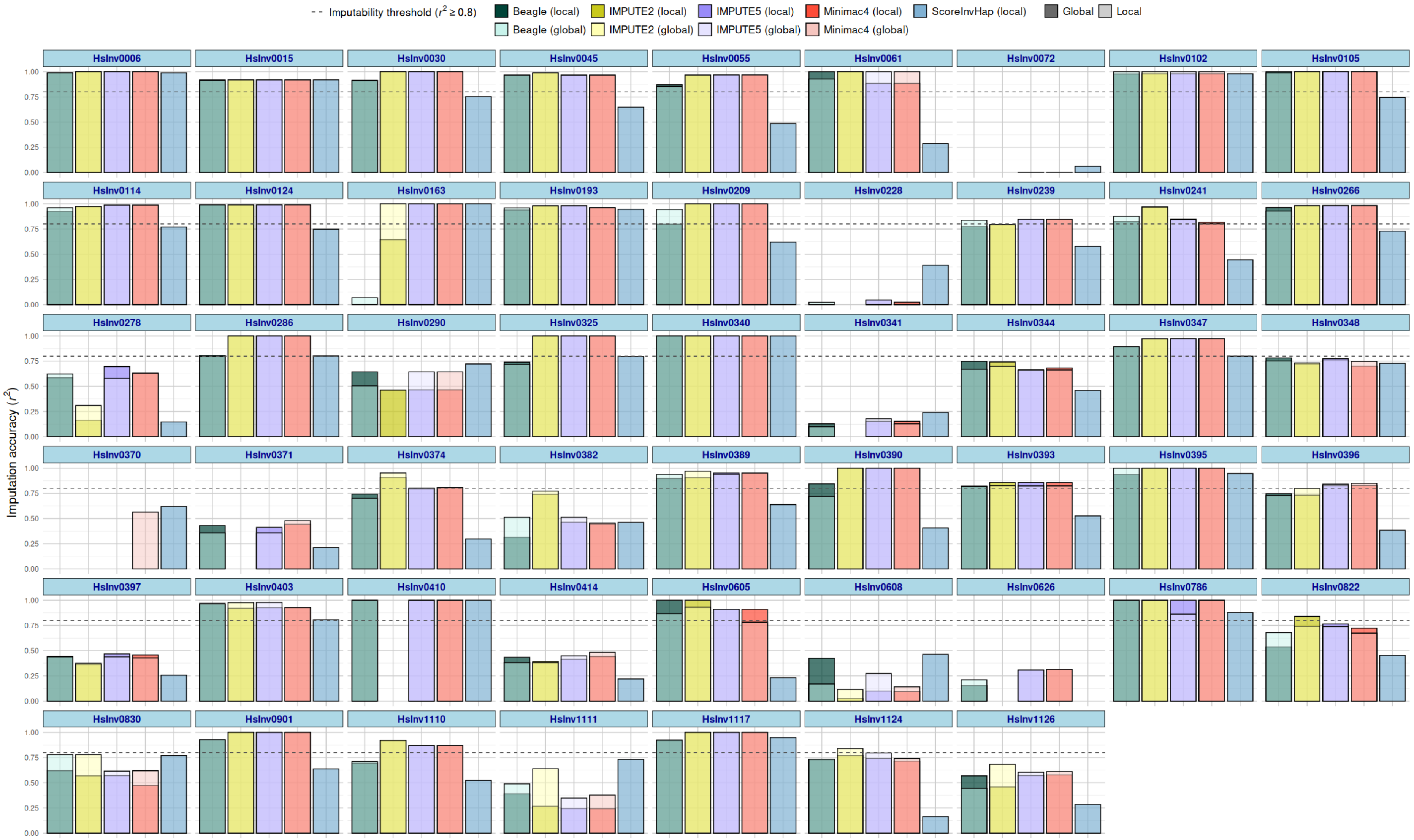

C

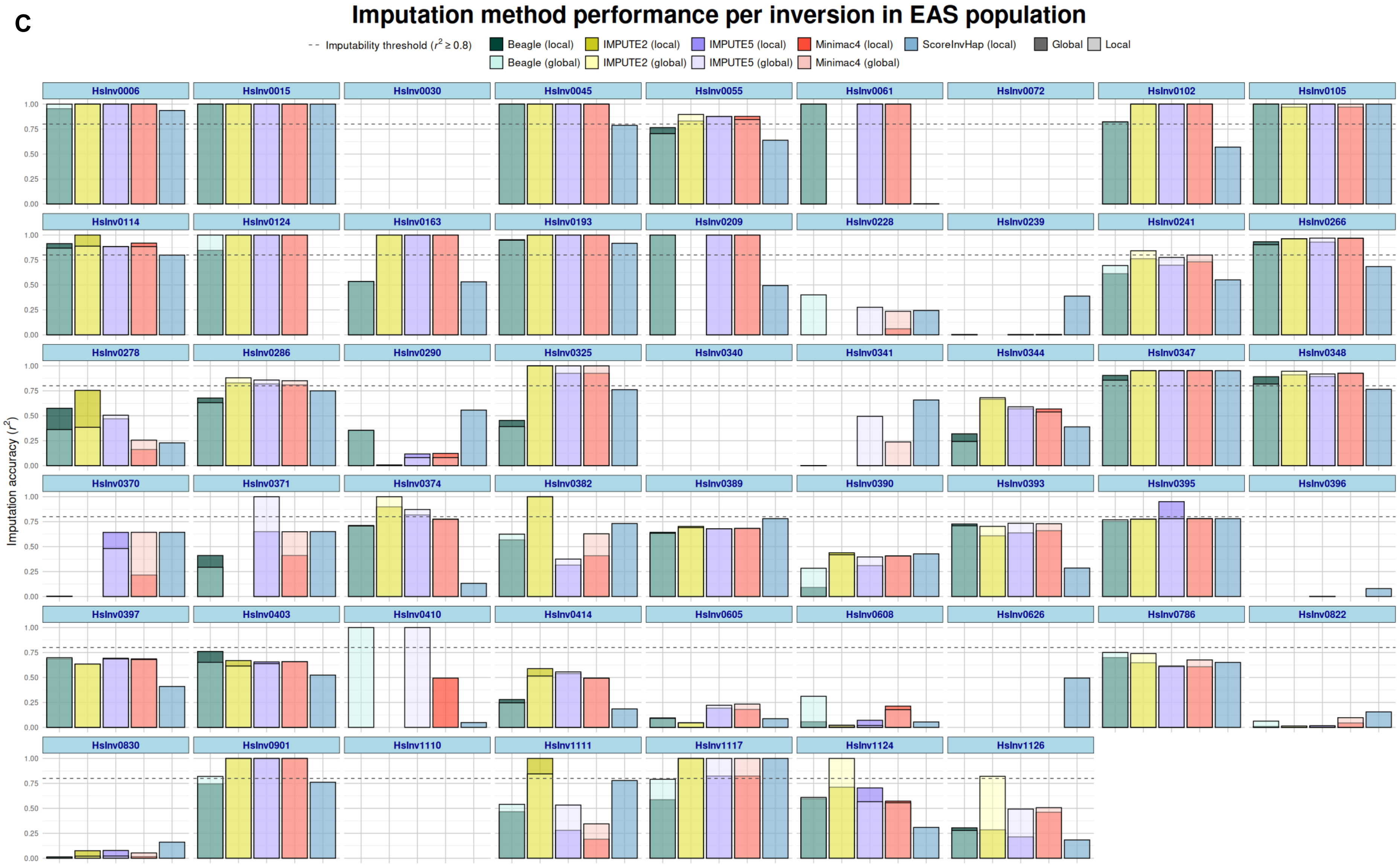

Imputation method performance per inversion in EAS population
Imputability threshold ($r^2 \geq 0.8$)
Beagle (local)
IMPUTE2 (local)
IMPUTE5 (local)
Minimac4 (local)
ScoreInvHap (local)
Global
Local
Beagle (global)
IMPUTE2 (global)
IMPUTE5 (global)
Minimac4 (global)
Imputation accuracy ($r^2$)
HsInv0006
HsInv0015
HsInv0030
HsInv0045
HsInv0055
HsInv0061
HsInv0072
HsInv0102
HsInv0105
HsInv0114
HsInv0124
HsInv0163
HsInv0193
HsInv0209
HsInv0228
HsInv0239
HsInv0241
HsInv0266
HsInv0278
HsInv0286
HsInv0290
HsInv0325
HsInv0340
HsInv0341
HsInv0344
HsInv0347
HsInv0348
HsInv0370
HsInv0371
HsInv0374
HsInv0382
HsInv0389
HsInv0390
HsInv0393
HsInv0395
HsInv0396
HsInv0397
HsInv0403
HsInv0410
HsInv0414
HsInv0605
HsInv0608
HsInv0626
HsInv0786
HsInv0822
HsInv0830
HsInv0901
HsInv1110
HsInv1111
HsInv1117
HsInv1124
HsInv1126
0.00
0.25
0.50
0.75
1.00

**Supplementary Figure 6.** Individual inversion imputation results in the three studied populations. Graphs show the imputation accuracy ($r^2$) of each inversion with the five tested programs (GP ≥ 0.9 filtering, except for scoreInvHap) using a population specific (local, dark color) or global (light color) reference panel in African (AFR, **A**), European (EUR, **B**) and East-Asian (EAS, **C**) populations. Dashed line indicates the $r^2 \geq 0.8$ threshold to consider an inversion as imputable.
